# Protein Fold Determination by Assembling Extended Super-Secondary Structure Motifs Using Limited NMR Data

**DOI:** 10.1101/509356

**Authors:** Kala Bharath Pilla

## Abstract

3D fold determination of proteins by computational algorithms guided by experimental restraints is a reliable and efficient approach. However, the current algorithms struggle with sampling conformational space and scaling in performance with increasing size of proteins. This paper demonstrates a new data-driven, time-efficient, heuristics algorithm that assembles the 3D structure of a protein from its elemental super-secondary structure motifs (Smotifs) using a limited number of nuclear magnetic resonance (NMR) derived restraints. The DINGO-NOE-RDC algorithm (3D assembly of Individual smotifs to Near-native Geometry as Orchestrated by limited nuclear Overhauser effects (NOE) and residual dipolar couplings (RDC)) leverages on the distance restraints recorded on methyl-methyl (CH_3_-CH_3_), methyl-amide (CH_3_-H^N^), and amide-amide (H^N^– H^N^) NOE contacts, and orientation restraints recorded via RDC on the backbone amide protons, to assemble the target’s Smotifs. Two conceptual advancements were made to bootstrap the structure determination from limited NMR restraints: Firstly, the basic definition of a ‘Smotif’ was expanded and secondly, a data driven approach for selection, scoring, ranking and clustering of Smotif assemblies is employed. In contrast to existing methods, the DINGO-NOE-RDC algorithm does not use a force-field or physical/empirical scoring function. Additionally, the algorithm employs a universal Smotif library that applies to any target protein and, can generate numerically reproducible results. For a benchmark set of ten different targets with different topologies, ranging from 100-200 residues, the algorithm identified near-native Smotifs for all of them.

## 1. Introduction

The accuracy of a protein structure can be assessed on two factors: Firstly, on the accuracy of the inter-atomic distances and secondly, the orientation of bond vectors relative to one another. Solution NMR spectroscopy can directly measure distance and bond orientation information in a target protein at near physiological conditions. 3D structure calculation using distance restraints rely on a substantial number of NOEs to provide a dense network of short-range distance restraints, usually, pairs of NMR active spins separated by distances up to 6 Å (Clore and Gronenborn, 1991; Wuthrich, 1986). Similarly, for protein molecules partially aligned to the magnetic field, the large one-bond internuclear dipolar interactions average to a non-zero value, resulting in observable residual dipolar coupling (RDC). The RDCs are precise reporters on the average orientation of bonds relative to the molecular alignment frame. *De novo* structure determination solely on the basis of RDCs is difficult due to the directional degeneracy inherent within the alignment tensor, which results in multiple local minima hindering the successful search for the native conformation (Bax, 2003). The greatest benefit of RDCs is in employing them to refine the orientation of bond vectors to an already determined atomic coordinate ensemble from NOEs. The resultant RDC refined structures present an ensemble that is precise in both inter-atomic distances and bond orientations. (Lipsitz and Tjandra, 2004).

In practice, the dense network of NOEs are difficult to resolve and with the increase in the size of the protein, the NOESY spectrum becomes increasingly complex resulting in resonance overlap. Additionally, slow tumbling of large proteins results in broadening of resonances thereby decreasing the sensitivity and resolution. In such cases, uniform deuteration of protein samples with selective labeling of methyl groups of isoleucine, leucine, valine and alanine (ILVA) allows identification of networks of methyl-methyl (CH_3_-CH_3_) NOEs (Ayala et al., 2009; Rosenzweig and Kay, 2014; Tugarinov and Kay, 2005). Combined with back-exchange of sidechain and amide protons with H_2_O, it enables additional identification of methyl-amide (CH_3_-H^N^), and amide-amide (H^N^–H^N^) NOE contacts (Tugarinov et al., 2006). While protein structure determination from just methyl-methyl and methyl-amide NOEs is difficult, protein folds of up to 80 kDa were determined by combining sparse NOE with multiple datasets of RDCs and employing rigid secondary structure elements (SSEs), torsion angle restraints and cartesian dynamics simulations (Mueller et al., 2000; Tugarinov et al., 2005).

ILVA-derived NOEs and RDCs were also utilized as restraints in hybrid structure prediction based on the iterative CS-RDC-NOE-Rosetta protocol, where protein domains up to 25 kDa were successfully determined (Lange et al., 2012; Raman et al., 2010). Rosetta follows a Monte-Carlo based fragment assembly approach to build 3D models of the target and the addition of NMR derived restraints not only help in steering the conformational search towards the native structure, which is otherwise intractable for large protein domains, but also aid in convergence (Baker, 2014; Fleishman and Baker, 2012; Pilla et al., 2016). A drawback of this approach is the usage of short fragment (three/nine residues) libraries which are not universal, but they are specific for the primary amino acid sequence of the target protein. The generated fragment libraries are also limited in number, in the case of Rosetta, up to 200 fragments can be used at each residue position. These restriction in fragment library size and generation is a compromise between minimizing conformational search space and maximizing the probability that the native protein structure can be accurately assembled from that fragments.

Structural motifs characterized as a group of SSEs connected by loops, such as Greek key, zinc finger, helix-turn-helix motifs, etc., are found in many protein families and are an attractive alternative to short fragments. The diversity of modern proteins and recurrence of these structural motifs is believed to reflect duplications, mutations, shuffling and fusion of genes through the course of evolution (Alva et al., 2015; Alva and Lupas, 2018; Lupas et al., 2001). These structural motifs represent a more natural description of the building blocks of proteins than an arbitrary choice of peptide fragment lengths. The elemental definition of a structural motif is two SSEs connected by a loop, called a Smotif (Fernandez-Fuentes et al., 2010). By this definition, there are only four basic types of Smotifs, which can be referred to as α-α, β-β, α-β and β-α, where α represents an alpha-helical element, and β represents extended polypeptide strand. Importantly, the total number of different Smotifs observed in all protein structures known to date has not changed since 2000 (last reported in 2010) (Fernandez-Fuentes et al., 2010), suggesting that our structural knowledge of Smotifs is likely to be complete. It was also shown that all known protein structures could be reconstructed with good accuracy from the finite set of Smotifs (Fernandez-Fuentes et al., 2010). Furthermore, Smotifs were employed in building topology-independent structure classification tools that are capable of quantitatively identifying the structural relationship between different topologies (Dybas and Fiser, 2016).

The quantitatively finite nature of Smotifs also predicts that a protein structure can be calculated if one can identify and assemble the target Smotifs. Previous attempts in using this approach were by two different methods. The first approach was by utilizing target-specific construction of Smotif libraries akin to Rosetta’s fragment library. Two demonstrations of this approach were, Smotifs in template-free modeling (SmotifTF) (Vallat et al., 2015) and chemical shift-guided Smotif assembly (SmotifCS) (Menon et al., 2013). Both algorithms performed on par with current state-of-the-art software like I-Tasser (Roy et al., 2010) and Rosetta (Rohl et al., 2004; Shen et al., 2009). A possible drawback of these approaches lies in the Monte Carlo sampling of protein structures, which limits their successful application to small proteins of around 110 residues. The second approach is in utilizing the universal Smotif libraries that are independent of any target. This was demonstrated using pseudocontact shift (PCS) NMR data, and it is the first implementation of DINGO-PCS algorithm (3D assembly of Individual Smotifs to Near-native Geometry as Orchestrated by PCSs) (Pilla et al., 2017). The DINGO-PCS utilizes PCSs of backbone amide protons as the only experimental data to identify, orient, and build the protein structure from its constituent Smotifs. The DINGO-PCS algorithm requires PCS datasets to be available from at least three different metal centers that are site-specifically incorporated in the target protein using artificial metal carrying chemical tags (Nitsche and Otting, 2017). The locations of these metal centers are ideally well dispersed in space to reduce correlation between PCS data sets, which can be difficult to achieve without prior knowledge of the target structure. Compared to PCSs, RDCs and NOEs are more difficult to measure but does not necessarily require any chemical modification of the target protein.

In this work, two major conceptual advancements are introduced. Firstly, the basic definition of Smotif is expanded from being a consecutive pair of SSEs connected by a loop to any pair of SSEs irrespective of the loop. Secondly, a data driven approach is employed for selection, scoring, ranking and clustering of Smotif assemblies, since the DINGO-NOE-RDC algorithm does not use a physical, empirical or statistical scoring function. The algorithm was benchmarked against ten proteins ranging from 100-200 residues in length and with different fold architectures, including membrane-bound, α-helical, β-barrel, and α/β topologies. All the ten chosen targets have experimental datasets recorded and published by different research groups and serve as a realistic benchmark set.

## 2. Results

The DINGO-NOE-RDC algorithm was first tested on Target-A (see Table-1) using a standard Smotif definition. An extended Smotif definition was implemented to test Target-B. Finally, the new extended Smotif definition was used to benchmark the performance of the algorithm on an additional set of eight proteins.

**Table 1:** Performance benchmark of the DINGO-NOE-RDC algorithm

| # | Targets | PDBID | $N_{\text{res}}^a$ | NOEs<br>used/available | $C^\alpha$<br>RMSD<sup>c</sup> | RDC<br>datasets | Sequence<br>identity<sup>d</sup> | BMRB<br>ID |
| --- | --- | --- | --- | --- | --- | --- | --- | --- |
| A | OmpX | 2MNH | 148 | 41/58 | 2.0 Å | 2 | 4.7% | 18796 |
| B | Human Prolactin | 1RW5 | 199 | 86/103 | 3.5 Å | 1 | 6.0% | 5599 |
| C | srr115c | 2KCV | 100 | 167/175 | 3.5 Å | 1 | 7.0% | 16084 |
| D | At5g22580 | 1RJJ | 111 | 66/73 | 3.7 Å | 1 | 8.1% | 6011 |
| E | Human MOG1 | 5YFG | 194 | 88/133 | 3.8 Å | 1 | 3.2% | 36117 |
| F | *De novo* designed<br>protein | 2LCI | 134 | 157/163 | 3.8 Å | 2 | 12.0% | 17613 |
| G | SgR145 | 2N47 | 202 | 188/385 | 4.5 Å | 1 | 5.5% | 16806 |
| H | Decorbin binding<br>protein | 2LQU | 168 | 129/145 | 4.6 Å | 1 | 5.3% | 18329 |
| I | ChR145 | 2LEQ | 146 | 140/182 | 4.9 Å | 3 | 6.8% | 17723 |
| J | ApaG | 2F1E | 127 | 47/62 | 5.2 Å | 1 | 5.2% | 5998 |

<sup>a</sup> Number of amino acid residues.
<sup>b</sup> the total number of available NOEs are counted only for methyl-methyl ( $\text{CH}_3\text{-CH}_3$ ), methyl-amide ( $\text{CH}_3\text{-H}^N$ ), and amide-amide ( $\text{H}^N\text{-H}^N$ ) NOEs for the entire chain of the target. The used number of NOEs are the total number that were utilized for the assembly of Smotifs.
<sup>c</sup> the $C^\alpha$ RMSD was calculated between the best Smotif assembly calculated by DINGO-NOE-RDC, which was identified as the structure best fulfilling the NOE and RDC data, and the Smotif residues in the corresponding reference structure.
<sup>d</sup> the sequence identity is computed by comparing the concatenated sequences from the assembled Smotifs to the native sequence make-up of the target.

### 2.1 DINGO-NOE-RDC on a ß-barrel integral membrane protein OmpX

The DINGO-NOE-RDC algorithm was first highlighted on an eight stranded anti-parallel ß-barrel fold, Outer membrane protein X (OmpX) from *Escherichia coli* (Hagn et al., 2013). The final assembly of all seven Smotifs was selected based on the experimental data that has the combined score for the largest number satisfied NOEs with the lowest distance violations and the lowest possible RDC fit score, see method section for more details. The final Smotif assembled structure as selected by the DINGO-NOE-RDC algorithm has a C^α^ RMSD of 2.0 Å when compared to the native NMR structure [PDBID: 2MNH] (Figure-1A; RMSD values were calculated for the residues covered by Smotifs only, unless explicitly stated otherwise). This is a notable result, considering that only 40 NOEs were used in computing the structure. The various parent structures from which the Smotifs were extracted are, 3D9R (ketosteroid isomerase-like protein from *Pectobacterium atrosepticum*), 1K24 (outer membrane adhesin from *Neisseria meningitidis*), 2VLG (KinA PAS-A domain from *Bacillus subtilis*), 1TUH (Bal32a from a soil-derived mobile gene cassette), 1JN5 (TAP-p15 mRNA export factor from *Homo sapiens*), 3Q7E (arginine methyltransferase from Rattus norvegicus), 4NZJ chain A (a putative galactosidase from *Bacteroides fragilis*). The sequence identity of the assembled Smotifs’ sequence compared to the target’s sequence is mere 4.7 % (Figure-1C) suggesting that the parent proteins are neither sequence nor structural homologs to Target-A. The DINGO-NOE-RDC algorithm does not use the target’s or the evaluating Smotifs’ sequence at any stage of assembly, and the low sequence identity reflects this. Additionally, the algorithm excludes native and near-native homologs from the Smotif assembly. OmpX and AIL (attachment invasion locus protein from *Yersinia pestis*, PDBID 3QRA), are currently the only experimentally resolved structures with eight anti-parallel ß-barrel fold, making it a target with a unique fold architecture. Moreover, except for 1K24, none of the parent proteins are membrane-bound, which shows that the DINGO-NOE-RDC algorithm can identify, select and assemble a target’s near-native Smotifs from the experimental input restraints only.

**Figure 1:**
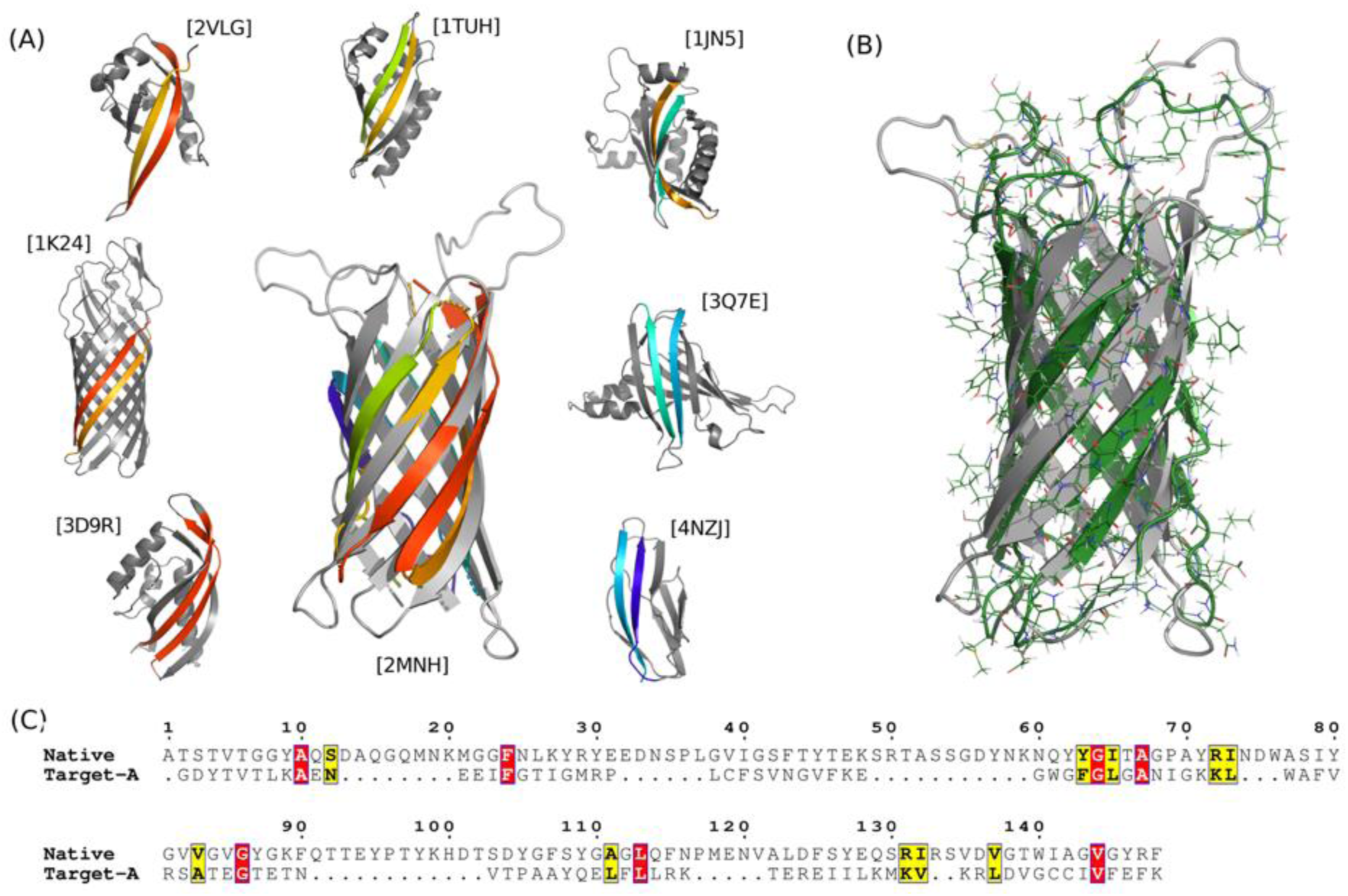
The Smotif assembly performed by DINGO-NOE-RDC algorithm on the integral membrane protein OmpX. (A) Cartoon representations of the final model superimposed onto the crystal structure (in gray) and of the parent proteins from which the Smotifs were derived. The respective parent proteins are labeled with their PDBID. The corresponding coloring highlights the Smotifs of the parent proteins and in the final structure. (B) All-atom model of the Smotif assembled structure from (A) generated using comparative modeling algorithm. (C) Sequence alignment between the target’s native sequence and assembled Smotifs is shown. Target-A’s sequence was generated by concatenating the individual sequences from the assembled Smotifs. Identical amino acid residues are highlighted in red and similar residues are highlighted in yellow. The sequence alignment is generated by placing equivalent residues between the target and the assembled Smotifs through secondary structure assignment.

Smotif assembly generates coordinates of the backbone and all-atom structures are easily computed by treating the final Smotif assembled model as the template and passing it as input to a comparative modeling algorithm; see Methods section on computing all-atom structures. The all-atom structure has a C^α^ RMSD of 4.0 Å, computed over all of the residues, to the reference NMR structure [PDBID: 2MNH], shown in Figure-1B.

### 2.2 Extending the Smotif definition

The standard definition of Smotif was originally proposed as a pair of SSEs that are consecutive, overlapping and connected by the loop for a given protein (Fernandez-Fuentes et al., 2010). This description of Smotifs was sufficient to determine the structure using a large number of pseudocontact shift restraints (Pilla et al., 2017). However, the standard Smotif definition was inefficient with the limited number of NOE and RDC data as demonstrated using Target-B (Table-1, Figure-2). Assigning Smotifs in the target by the standard definition, the DINGO-NOE-RDC algorithm was able to assemble only three out of four Smotifs in the Target-B, Figure-2B.

**Figure 2:**
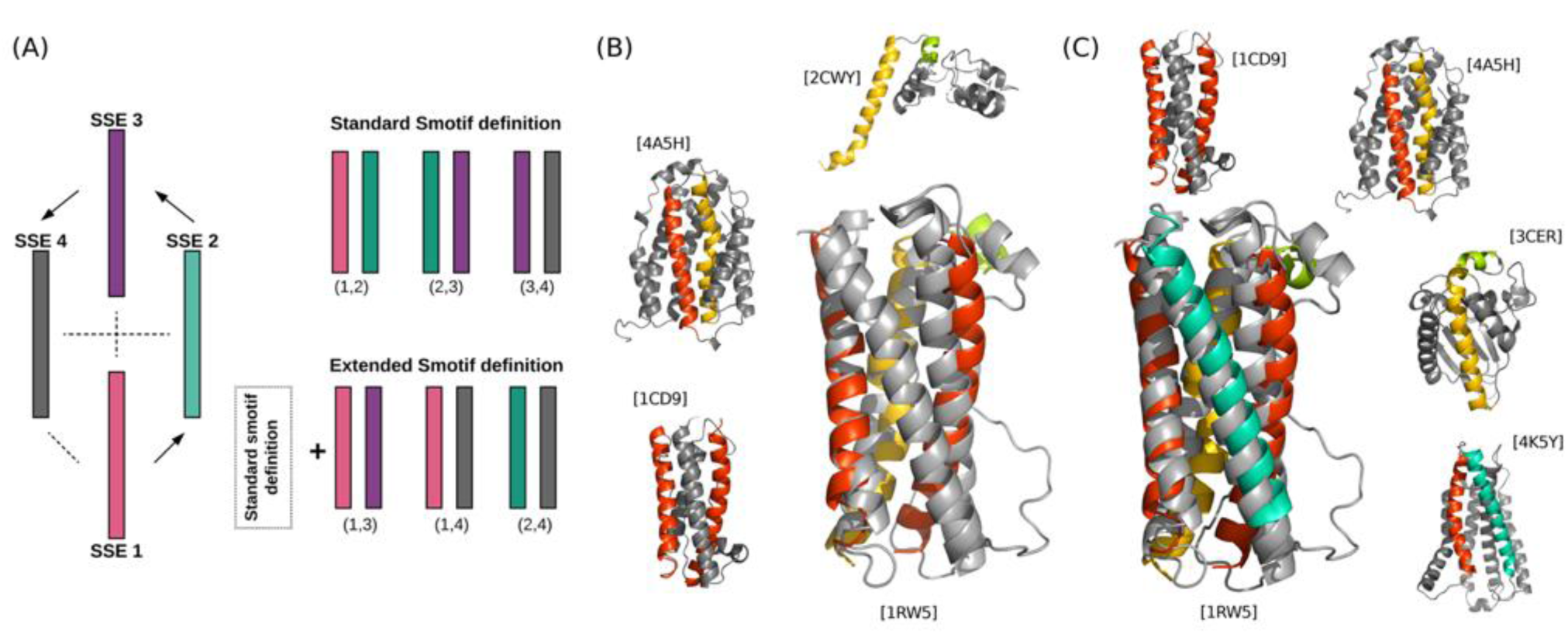
(A) Schematic representation depicting the extension of the Smotif definition on a four-helix bundle. The standard and extended definitions spanning various helical pairs are shown. (B) Target-B’s Smotif assembly by only considering the standard Smotif definition. The cartoon representations of the final model superimposed onto the crystal structure (in gray) and of the parent proteins from which the Smotifs were derived. The respective parent proteins are labeled with their PDBID. The Smotifs of the parent proteins and in the final structure is highlighted by the corresponding coloring. (C) Same as (B) but the extended definition of the Smotifs are considered.

The standard Smotif definition can be extended from sequential overlapping pairs of a given protein to all possible pairs of SSEs if one ignores the interconnecting loop. On a positive effect, this extended definition also increases the conformational search space encoded within the Smotifs. However, the number of Smotifs assignments also increases combinatorially with the number of SSEs in a given target. To minimize this impact, the DINGO-NOE-RDC algorithm considers all possible pairs of Smotifs only at a given stage in the assembly path. I.e., for the first pair of Smotifs, no extended Smotif definitions were considered, but for the subsequent Smotif, the possible Smotif definitions with the third SSE is considered. For example, consider a target with four SSEs. The first Smotif pair is (SSE-1, SSE-2) and no extended Smotifs are considered, but for the next step in assembly, both standard Smotif definition from overlapping SSE pairs (SSE-2, SSE-3) and the extended Smotif definition (SSE-1, SSE-3) are considered. For the final Smotif, the standard Smotif definition is (SSE-3, SSE-4) and the extended Smotif definitions does additionally include (SSE-1, SSE-4) and (SSE-2, SSE-4), shown in Figure-2A. Out of all of the extended Smotifs, the Smotif that best satisfies the experimental data is retained while discarding the poor fitting Smotif. This extended definition of Smotifs was essential for the successful assembly of Smotifs for Target-B which consists of four Smotifs, Figure-2C. In case of Target-B, the first three Smotifs were assembled from sequential assignment of Smotif definition (Figure-2B), but for the fourth Smotif, the extended Smotif definition considering the SSE pair (SSE-1, SSE-5) was needed, as the sequential Smotif assignment pair (SSE-4, SSE-5) yielded no Smotif that could satisfy the experimental data, see Figure-2C.

### 2.3 DINGO-NOE-RDC Performance Benchmark

The DINGO-NOE-RDC was benchmarked on an additional set of eight targets (Table-1). The benchmark set was chosen to include targets with different topologies. All of the NOE and RDC datasets used in the evaluation of the DINGO-NOE-RDC algorithm were previously published, and their reference BMRB IDs are listed in Table-1. No synthetic datasets were used. If the experiment dataset contains NOEs for spins other than ILVA (methyl) and backbone-amide protons, the additional NOEs were discarded to reduce the data. The quality of assembled Smotifs ranged from 2.0 Å – 5.2 Å C^α^ RMSD when compared to the native NMR structures. The superpositions of the computed structures with their corresponding reference structures are shown in Figure-3. The C^α^ RMSD was computed against the first model of the ensemble when more than one structure was reported. The sequence identity from the assembled Smotifs when compared to the native sequence make-up of the target is below 12.0 %, implying that the DINGO-NOE-RDC algorithm selected no native, near-native or structural homologs. Surprisingly, the highest sequence identity was found for the Target-F, which was claimed to be a *de novo* designed protein [PDBID:2LCI].

**Figure 3:**
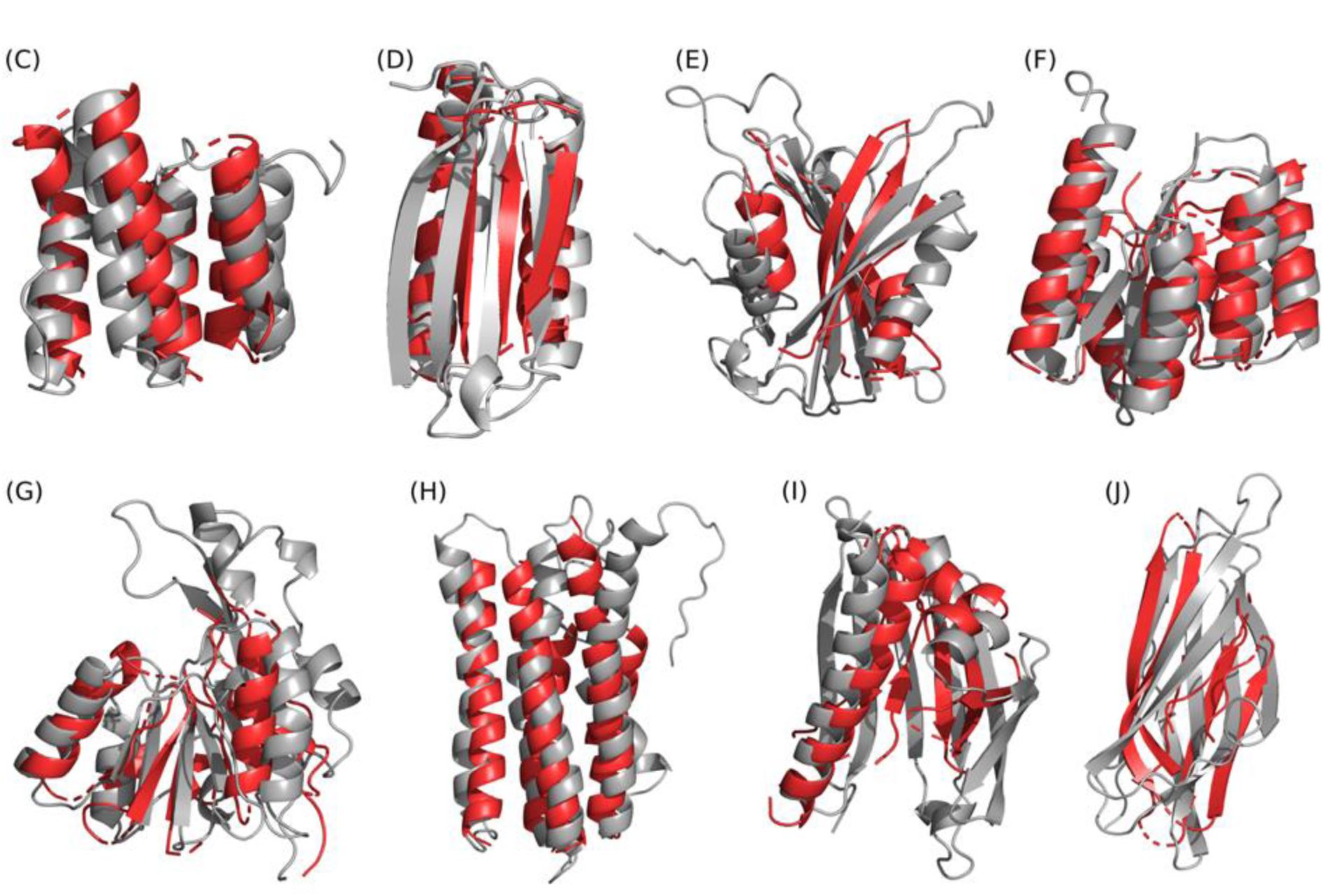
Superpositions of the best Smotif assemblies for Targets (C-J) calculated with DINGO-NOE-RDC (red) onto the corresponding reference structures (gray). The best Smotif assemblies were identified as the assemblies best satisfying the experimental NOE and RDC data. The targets are labeled as in Table-1 (See also Figures S1-S8).

## 3. Discussion

The DINGO-NOE-RDC method is distinct in its approach to explore the conformational space associated with the target sequence, especially, when compared to Rosetta or molecular dynamics methods. Rosetta relies on sequence assisted selection of fragment libraries and a physical and knowledge-based scoring function to explore the conformational space (Alford et al., 2017), and molecular dynamics rely on the physical principles encoded via the force-fields to explore the conformational space in a time-dependent manner (Brooks et al., 2009). While both Rosetta and molecular dynamics scoring functions perform on par with one another (Rubenstein et al., 2018), The DINGO approach does not use neither of these approaches but exploits the fixed conformational space encoded within the Smotifs. The DINGO approach is a genetic algorithm and does not have any random sampling routines which makes the Smotif assemblies of any target repeatable and numerically reproducible.

The implementation of extended Smotif definition was essential for the successful assembly for nine out of ten targets. The Smotif assembly of Target-A was the only structure determined using the standard Smotif definition. The use of the extended Smotif definition for this target did not change the outcome of the final Smotif assembly. The sparsity of the input restraints and the DINGO-NOE-RDC algorithm being dependent only on the input data, degenerate solutions were observed for Target-E, shown in Figure-S9. The C terminal helix, spanning the residues 164 to 174, was not correctly positioned among similar scoring top Smotif assemblies, which is because no NOEs are being observed between the C terminal helix and the other SSEs. Although RDCs were available, the dataset from a single alignment tensor was not sufficient for the algorithm to accurately position the C terminal helix, resulting in degenerate solutions.

The DINGO-NOE-RDC algorithm can identify structural features purely from the experimental input data, demonstrated with Target-C’s tetratricopeptide repeat (TPR) domain. TPR is a 34-amino acid α-helical motif spanning two antiparallel α-helices and found in over 300 different proteins (Blatch and Lässle, 1999; Main et al., 2003). Target-C [PDBID:2KCV] contains two TPR repeats, covering over four N terminal α-helices. The DINGO-NOE-RDC algorithm assembled the first three of the four TPR repeat domains by repeatedly assembling a single Smotif, shown in Figure-S1. The fourth helix is not covered by the repeat Smotif, as the native helix is of a different length than the first TPR domain.

The computational cost associated with the exhaustive search of Smotif libraries is low. For example, Target-A took only 50 CPU hours on an Intel Xeon Sandy Bridge 2.6 GHz processor to compute the Smotifs. The only limiting step in the exhaustive enumeration of Smotifs is the quantity of Smotifs binned to the definition of Smotif. For Target-B, which contains four large α-helical Smotifs spanning 199 residues, it took 75 CPU hours for enumeration of extended Smotifs. However, for Target-G spanning 202 residues consisted of 11 short Smotif definitions it took ~2500 CPU hours or ~20 hours on a 128 CPU cluster. The DINGO-NOE-RDC algorithm is implemented to take advantage of parallel processing and can be scaled nearly linear with the number of processors. In contrast, Rosetta requires between 5 and 100 times more computational power for similar sized proteins and with similar experimental restraints (Lange et al., 2012; Lange and Baker, 2012; Raman et al., 2010).

The input NMR datasets used primarily contained NOE restraints from ILVA methyl sidechains and methyl-amides. This is because these restraints provide the most long-range distance information and can be measured on large protein systems. DINGO-NOE-RDC algorithm can utilize any NOEs that are recorded between any pairs of protons. Additionally, this algorithm can be easily customized to score a diverse range of restraints, like PCSs, and distance restraints from paramagnetic relaxation enhancements (PRE) and electron paramagnetic resonance (EPR).

The DINGO-NOE-RDC algorithm uses experimental data in every step of scoring, selection, and ranking of modeled structures. In doing so, the NOE and RDC datasets are assumed to have no ambiguity associated with their assignments. Additionally, the conformational space of any target is assumed to be sufficiently represented in our Smotif libraries. While there was not any target in this benchmark set that has failed to assemble, but there was a case that was previously reported using PCSs where the DINGO-PCS failed to find any Smotif of a specific orientation within its Smotif libraries that resulted in incomplete Smotif assembly (Pilla et al., 2017).

The DINGO-NOE-RDC software package can be downloaded for free along with the results of the benchmark proteins from the Git hub repository https://github.com/kalabharath/DINGO_NOE_RDC, and the precompiled universal Smotif libraries are freely available to download from http://comp-bio.anu.edu.au/huber/Smotifs/

## 4. Conclusion

The DINGO-NOE-RDC algorithm presents a new way to determine 3D folds of proteins using sparse NOEs and RDCs as the only input. The success of this method relies on the inclusion of the extended definition of Smotifs. The DINGO-NOE-RDC algorithm performed consistently in calculating the 3D structure for all tested benchmark proteins, with RMSDs in the range of 2.0 Å - 5.2 Å compared to the reference NMR structures.

## 5. Methods

### 5.1. Generating universal Smotif library

A nonredundant Smotif library was generated using 308,999 domains obtained from the CATH database (Sillitoe et al., 2015). To extract and build the Smotif libraries, one needs only the secondary structure assignment of the domain and the coordinates of backbone atoms. The program STRIDE (Frishman and Argos, 1995) was used to assign the SSEs for all CATH domains. The SSEs were represented by ‘E’ for β-strands and ‘H’ for helices, including α-helices, 3_10_-helices, and Π-helices. Alanine residues were added to domain chain breaks less than ten residues long and to the missing electron density for residues within SSEs to avoid distortion of smotif assignment. 3D coordinates of the added alanine residues were generated using the comparative modeling package Modeller (Webb and Sali, 2014). For domains with larger than ten residues chain breaks, the SSEs flanking the chain breaks were not assigned to any Smotif definition. For CATH domains from X-ray crystallographic structures, the backbone amide hydrogens were added using ‘pdb2gmx’ program from the GROMACS software package (Van Der Spoel et al., 2005). To avoid the loss of backbone amide hydrogen data from the Proline residues, the Smotifs were modified by adding a pseudo-hydrogen to the backbone amide.

The universal library consisted of Smotifs where each SSE was at least five residues in length and devoid of loops. Smotifs with two β-strands SSEs consisting of 10 and 5 residues were added to a file labeled as ‘ss_10_5.db’. Similarly, other Smotifs were sorted into individual files based on the length of their respective SSEs. The Smotifs in each file are further clustered with a radius of 0.25 Å to one another so as to remove structurally redundant entries. The new version of the Smotif library has 555,558 entries grouped in 3,405 Smotif entries, which is 27 % more entries than the previous database built on CATH’s S100 sequence nonredundant release (Pilla et al., 2017).

### 5.2. Secondary structure assignment of the target protein

TALOS-N server generates the secondary structure assignments for the target proteins using backbone chemical shifts to predict torsion angles and assigns secondary structure with an accuracy of 89 % (Shen and Bax, 2013). To allow for errors in the secondary structure assignment, each SSE is allowed to vary by 11 different permutations [(1, 0), (2, 0), (−1, 0), (−2, 0), (0, 1), (0, 2), (0, −1),(0, −2), (−1, −1),(1, 1), and (0, 0)], where the first and second numbers give the addition/truncation of residues at the N and C termini of the SSE.

### 5.3. Computing sidechains for methyl labeled amino acids

Backbone-dependent sidechain rotamer libraries were generated for each of the ILVA residues using a two-step approach.

1. In the first step, the various backbone conformations of the ILVA residues were established from a randomly selected 50,000 protein domains. The conformations were binned into different clusters with an arbitrary cutoff 0.1 Å RMSD radius from one another. For example, isoleucine has 142 different bins of backbone conformations.
2. In each of the back-bone clustered bins, the side chains were further clustered at a 0.5 Å RMSD radius from one another. For example, the 142 different backbone conformations of the Isoleucine consist of 2,159 discrete side-chain rotamers.

### 5.4. 3D assembly of Individual Smotifs to Near-native Geometry as Orchestrated by limited NOE and RDC data (DINGO-NOE-RDC)

The target protein’s secondary structure assignment was used to assign the Smotifs. The Smotifs were then ranked by the amount of RDC data each Smotif contains. A Smotif assembly sequence was then generated starting from the Smotif associated with the largest number of RDCs. The next overlapping Smotif was chosen based on the number of RDCs available for it, resulting in either N or C terminal extension of the initial Smotif. See the schematic flowchart in Figure 4 describing the various steps involved in the DINGO-NOE-RDC algorithm. To simplify the data guided Smotif assembly, a two-stage process was employed, (1) identifying the initial Smotif and (2) subsequently extending the following Smotifs.

**Figure 4:**
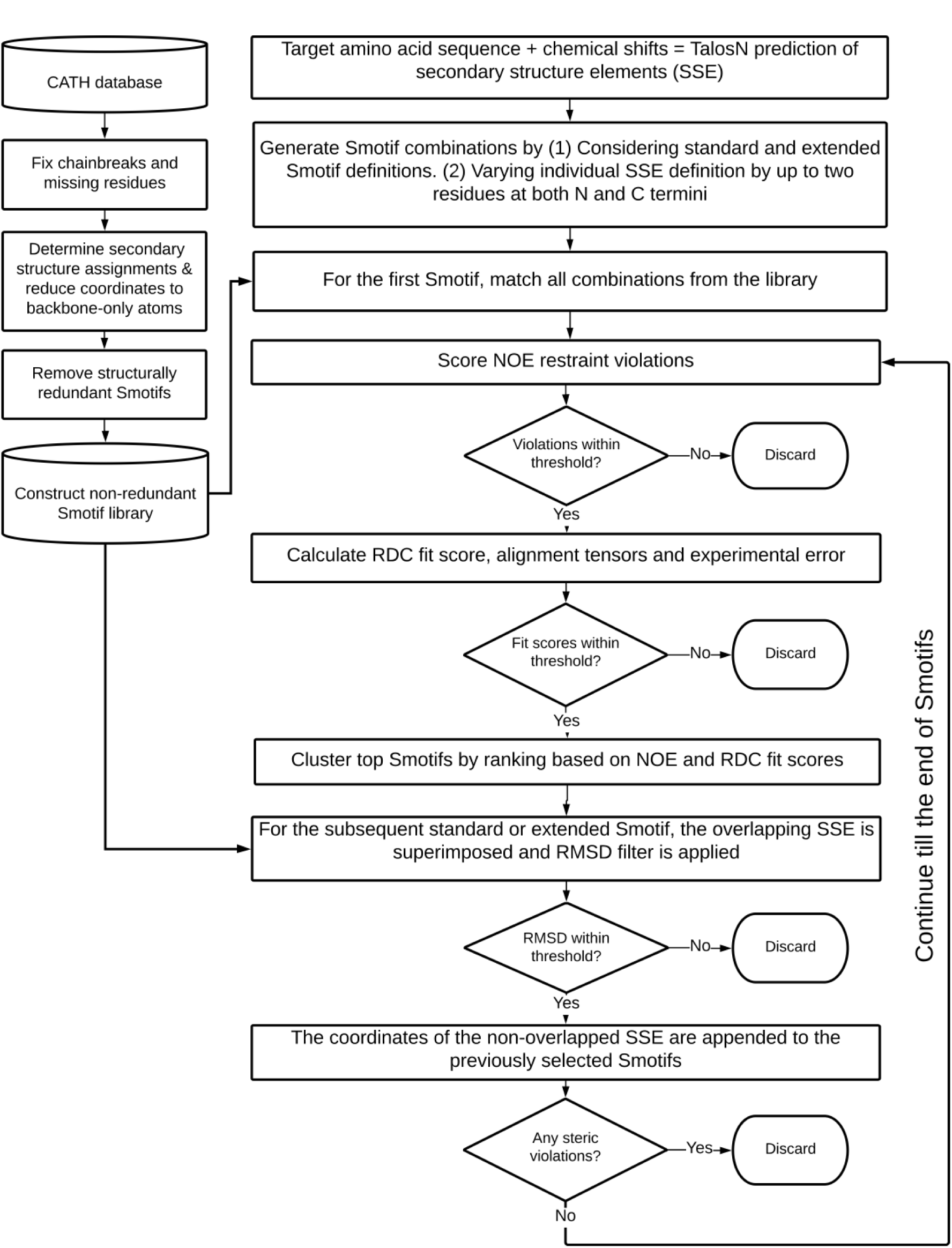
Flowchart describing the various steps involved in the DINGO-NOE-RDC algorithm.

#### 5.4.1. Stage-1: Identifying the initial Smotif

For the initial Smotif with two SSEs and accounting the 11 different permutation changes in the secondary structure assignment per SSE, it adds up to a total of 121 combinations to be evaluated. All of the Smotif entries in the 121 possible Smotif combinations were exhaustively searched. Each Smotif was evaluated against experimental data and it is considered a potential hit if it passes through the three data filters:

1. NOE filter: If the Smotif contains any back-calculated NOEs whose distance is greater than 6.0 Å, or less than 1.8 Å, it is discarded without any further valuation, else, the back-calculated NOEs must agree with the experimental value within an error threshold as defined by the experiment and the total NOE fit score is calculated as:

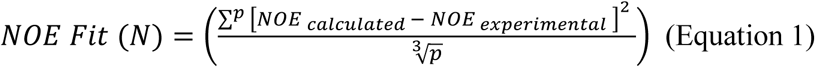

Where *p* is the total number of available NOEs.
2. RDC filter: Similar to the NOE filter, if any of the back-calculated RDC value exceeds the error threshold of 10.0 Hz, the Smotif is discarded without further evaluation, else, the back-calculated RDCs must agree with the experimental value within an error threshold as defined by the experiment and the total RDC fit score is calculated as:

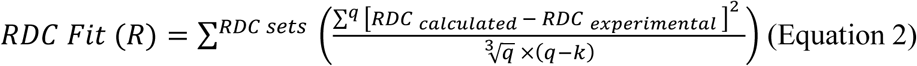

Where *q* is the total number of available RDCs per dataset and k is the total number of fitted parameters per dataset.
3. Alignment tensor filter: Additional filter is placed on the alignment tensor parameters. The magnitudes of the axial and rhombic components of the alignment tensors were restricted to a default value ±50 Hz. This value can be user modified. These boundaries aid in the filtering of incorrect Smotifs, especially when the Smotifs contain RDCs that have small magnitudes or, fewer in number, which can be easily fitted by unrealistic axial and rhombic parameters.

All the hits are ranked in a three-step process. In the first step, all the assemblies are binned according to their total number of satisfied NOEs and within each bin, the Smotif assemblies are ranked according to their NOE fit score (N), rounded to two decimal places, as described in equation (1). In the second step, for a given NOE fit score (N), all the assembles were further ranked based on their RDC fit score (R) as given by equation (2). In the third-step, the ranked assemblies were screened for structural redundancy (with an arbitrary cutoff of 1.0 Å) and only top 128 hits (or a user-defined number) are taken to the next stage. For this benchmark, only 128 Smotif assemblies were chosen for convenience of parallel scheduling calculation on a batch of 128 CPUs.

#### 5.4.2. Stage-2: Extension the Smotif assembly

In this stage both standard and extended Smotifs definitions were considered. The next Smotif can be added to either N or C termini of the initial Smotif based on the number of RDCs it contains. If the same number of RDCs characterizes the subsequent Smotif as the initial Smotif, the direction of Smotif assembly is decided by the number of RDCs available for the next subsequent Smotif. A five-step approach described below is followed to continue with the Smotif assembly.

Step 1: In this step, the length of the SSE shared between the previous Smotif and the current Smotif is fixed, while the other SSE in the current Smotif is varied by 11 different permutations as described in the stage-1.

Step 2: By definition, Smotifs come in overlapping pairs, and this enables us to filter out non-overlapping Smotifs without evaluating them for NOE and RDC fitting. A threshold RMSD, less than 2.0 Å is accepted for the overlapping SSE. The Newton-Raphson quaternion-based RMSD calculation algorithm (Liu et al., 2010) is used in this step. The algorithm was capable of calculating approximately 7000 RMSDs per second, enabling rapid filtering of Smotifs. The remaining Smotifs were taken to the next step.

Step 3: In this step, the newly identified Smotif coordinates were translated to the coordinate frame of the previous Smotif and the coordinates of the overlapping SSE in the current Smotif were discarded. The coordinates of the non-overlapping SSE were appended to the previous Smotif assembly. The assembly was further filtered for any backbone clashes with other SSEs. Those structures that pass this filter were further propagated to the next step.

Step 4: The three experimental data filters described in Stage 1 are reapplied, and the Smotif assemblies that satisfy all three filters are considered potential hits.

Step 5: In this final step, the potential hits were further filtered for structural redundancies. All of the hits were ranked according to the agreement with the experimental data and then propagated for the next round of selection of Smotifs. The process is repeated until all the Smotifs of the target protein has been assembled.

The DINGO-NOE-RDC is a genetic algorithm, where a population of Smotif assemblies was evaluated and propagated in each stage. If no Smotifs survive the filters in any given stage, the assembly is terminated and marked as incomplete. The completed Smotif assemblies were ranked based on their quality of NOE and RDC fit and the best-fitting Smotif and the Smotif assembly is reported as the final model. The NOE and RDC quality can vary dramatically from the first-ranked Smotif assembly to the second-ranked assembly. Therefore, only the best-ranked assembly was reported.

### 5.5. Generating all-atom models

As the Smotif assemblies compute only coordinates of backbone atoms and are devoid of loops, models need to be transformed into all-atom models by employing a comparative modeling software, treating the assembled model as structural template. Here, RosettaCM protocol was employed for generating all-atom models (Song et al., 2013). For each target 1000 models were generated. These models were further refined using the Rosetta Relax protocol (Conway et al., 2014), generating 10,000 models from the initial 1000 structures. The top model was selected based on the best fit to NOE and RDC data, ranked using Rosetta’s inbuilt NOE and RDC scoring functions (Raman et al., 2010).

## Supporting information

Supplemental Images

## Acknowledgments

K.B.P acknowledges Prof. Thomas Huber for his support with the computational resources, critical comments and feedback on the manuscript. K.B.P also acknowledges Dr. Nandhitha Subramanian for her comments and suggestions on the manuscript.

## References

Alford RF, Leaver-Fay A, Jeliazkov JR, O’Meara MJ, DiMaio FP, Park H, Shapovalov M V., Renfrew PD, Mulligan VK, Kappel K, Labonte JW, Pacella MS, Bonneau R, Bradley P, Dunbrack RL, Das R, Baker D, Kuhlman B, Kortemme T, Gray JJ. 2017. The Rosetta All-Atom Energy Function for Macromolecular Modeling and Design. J Chem Theory Comput 13:3031–3048. doi:10.1021/acs.jctc.7b00125

Alva V, Lupas AN. 2018. From ancestral peptides to designed proteins. Curr Opin Struct Biol 48:103–109. doi:10.1016/j.sbi.2017.11.006

Alva V, Söding J, Lupas AN. 2015. A vocabulary of ancient peptides at the origin of folded proteins. Elife 4:e09410. doi:10.7554/eLife.09410

Ayala I, Sounier R, Usé N, Gans P, Boisbouvier J. 2009. An efficient protocol for the complete incorporation of methyl-protonated alanine in perdeuterated protein. J Biomol NMR 43:111–119. doi:10.1007/s10858-008-9294-7

Baker D. 2014. Centenary Award and Sir Frederick Gowland Hopkins Memorial Lecture. Protein folding, structure prediction and design. Biochem Soc Trans 42:225–229. doi:10.1042/BST20130055

Bax A. 2003. Weak alignment offers new NMR opportunities to study protein structure and dynamics. Protein Sci 12:1–16. doi:10.1110/ps.0233303

Blatch GL, Lässle M. 1999. The tetratricopeptide repeat: a structural motif mediating protein-protein interactions. BioEssays 21:932–939. doi:10.1002/(SICI)1521-1878(199911)21:11<932::AID-BIES5>3.0.CO;2-N

Brooks BR, Brooks CL, Mackerell AD, Nilsson L, Petrella RJ, Roux B, Won Y, Archontis G, Bartels C, Boresch S, Caflisch A, Caves L, Cui Q, Dinner AR, Feig M, Fischer S, Gao J, Hodoscek M, Im W, Kuczera K, Lazaridis T, Ma J, Ovchinnikov V, Paci E, Pastor RW, Post CB, Pu JZ, Schaefer M, Tidor B, Venable RM, Woodcock HL, Wu X, Yang W, York DM, Karplus M. 2009. CHARMM: The biomolecular simulation program. J Comput Chem 30:1545–1614. doi:10.1002/jcc.21287

Clore GM, Gronenborn AM. 1991. Structures of larger proteins in solution: three-and four-dimensional heteronuclear NMR spectroscopy. Science (80-) 252:1390–1399. doi:10.1126/SCIENCE.2047852

Conway P, Tyka MD, DiMaio F, Konerding DE, Baker D. 2014. Relaxation of backbone bond geometry improves protein energy landscape modeling. Protein Sci 23:47–55. doi:10.1002/pro.2389

Dybas JM, Fiser A. 2016. Development of a motif-based topology-independent structure comparison method to identify evolutionarily related folds. Proteins Struct Funct Bioinforma 84:1859–1874. doi:10.1002/prot.25169

Fernandez-Fuentes N, Dybas JM, Fiser A. 2010. Structural characteristics of novel protein folds. PLoS Comput Biol 6:e1000750. doi:10.1371/journal.pcbi.1000750

Fleishman SJ, Baker D. 2012. Role of the biomolecular energy gap in protein design, structure, and evolution. Cell 149:262–273. doi:10.1016/j.cell.2012.03.016

Frishman D, Argos P. 1995. Knowledge-based protein secondary structure assignment. Proteins Struct Funct Bioinforma 23:566–579. doi:10.1002/prot.340230412

Hagn F, Etzkorn M, Raschle T, Wagner G. 2013. Optimized Phospholipid Bilayer Nanodiscs Facilitate High-Resolution Structure Determination of Membrane Proteins. J Am Chem Soc 135:1919–1925. doi:10.1021/ja310901f

Lange OF, Baker D. 2012. Resolution-adapted recombination of structural features significantly improves sampling in restraint-guided structure calculation. Proteins Struct Funct Bioinforma 80:884–895. doi:10.1002/prot.23245

Lange OF, Rossi P, Sgourakis NG, Song Y, Lee H-W, Aramini JM, Ertekin A, Xiao R, Acton TB, Montelione GT, Baker D. 2012. Determination of solution structures of proteins up to 40 kDa using CS-Rosetta with sparse NMR data from deuterated samples. Proc Natl Acad Sci 109:10873–10878. doi:10.1073/pnas.1203013109

Lipsitz RS, Tjandra N. 2004. Residual dipolar couplings in NMR structure analysis. Annu Rev Biophys Biomol Struct 33:387–413. doi:10.1146/annurev.biophys.33.110502.140306

Liu P, Agrafiotis DK, Theobald DL. 2010. Rapid communication fast determination of the optimal rotational matrix for macromolecular superpositions. J Comput Chem 31:1561– 1563. doi:10.1002/jcc.21439

Lupas a N, Ponting CP, Russell RB. 2001. On the evolution of protein folds: are similar motifs in different protein folds the result of convergence, insertion, or relics of an ancient peptide world? J Struct Biol 134:191–203. doi:10.1006/jsbi.2001.4393

Main ERG, Xiong Y, Cocco MJ, D’Andrea L, Regan L. 2003. Design of stable α-helical arrays from an idealized TPR motif. Structure 11:497–508. doi:10.1016/S0969-2126(03)00076-5

Menon V, Vallat BK, Dybas JM, Fiser A. 2013. Modeling Proteins Using a Super-Secondary Structure Library and NMR Chemical Shift Information. Structure 21:891–899. doi:10.1016/j.str.2013.04.012

Mueller GA, Choy WY, Yang D, Forman-Kay JD, Venters RA, Kay LE. 2000. Global folds of proteins with low densities of NOEs using residual dipolar couplings: Application to the 370-residue maltodextrin-binding protein. J Mol Biol 300:197–212. doi:10.1006/jmbi.2000.3842

Nitsche C, Otting G. 2017. Pseudocontact shifts in biomolecular NMR using paramagnetic metal tags. Prog Nucl Magn Reson Spectrosc 98–99:20–49. doi:10.1016/j.pnmrs.2016.11.001

Pilla KB, Otting G, Huber T. 2017. Protein structure determination by assembling super-secondary structure motifs using pseudocontact shifts. Structure 25:559–568. doi:10.1016/j.str.2017.01.011

Pilla KB, Otting G, Huber T. 2016. Pseudocontact Shift-Driven Iterative Resampling for 3D Structure Determinations of Large Proteins. J Mol Biol 428:522–532. doi:10.1016/j.jmb.2016.01.007

Raman S, Lange OF, Rossi P, Tyka M, Wang X, Aramini J, Liu G, Ramelot T a, Eletsky A, Szyperski T, Kennedy M a, Prestegard J, Montelione GT, Baker D. 2010. NMR Structure Determination for Larger Proteins Using Backbone-Only Data. Science (80-) 327:1014– 1018. doi:10.1126/science.1183649

Rohl CA, Strauss CEM, Misura KMS, Baker D. 2004. Protein structure prediction using Rosetta. Methods Enzymol 383:66–93. doi:10.1016/S0076-6879(04)83004-0

Rosenzweig R, Kay LE. 2014. Bringing Dynamic Molecular Machines into Focus by Methyl-TROSY NMR. Annu Rev Biochem 83:291–315. doi:10.1146/annurev-biochem-060713-035829

Roy A, Kucukural A, Zhang Y. 2010. I-TASSER: a unified platform for automated protein structure and function prediction. Nat Protoc 5:725–738. doi:10.1038/nprot.2010.5

Rubenstein AB, Blacklock K, Nguyen H, Case DA, Khare SD. 2018. Systematic Comparison of Amber and Rosetta Energy Functions for Protein Structure Evaluation. J Chem Theory Comput 14:6015–6025. doi:10.1021/acs.jctc.8b00303

Shen Y, Bax A. 2013. Protein backbone and sidechain torsion angles predicted from NMR chemical shifts using artificial neural networks. J Biomol NMR 56:227–241. doi:10.1007/s10858-013-9741-y

Shen Y, Vernon R, Baker D, Bax A. 2009. De novo protein structure generation from incomplete chemical shift assignments. J Biomol NMR 43:63–78. doi:10.1007/s10858-008-9288-5

Sillitoe I, Lewis TE, Cuff A, Das S, Ashford P, Dawson NL, Furnham N, Laskowski RA, Lee D, Lees JG, Lehtinen S, Studer RA, Thornton J, Orengo CA. 2015. CATH: comprehensive structural and functional annotations for genome sequences. Nucleic Acids Res 43:D376– D381. doi:10.1093/nar/gku947

Song Y, Dimaio F, Wang RY-R, Kim D, Miles C, Brunette T, Thompson J, Baker D. 2013. High-Resolution Comparative Modeling with RosettaCM. Structure 21:1735–1742. doi:10.1016/j.str.2013.08.005

Tugarinov V, Choy W-Y, Orekhov VY, Kay LE. 2005. Solution NMR-derived global fold of a monomeric 82-kDa enzyme. Proc Natl Acad Sci 102:622–627. doi:10.1073/pnas.0407792102

Tugarinov V, Kanelis V, Kay LE. 2006. Isotope labeling strategies for the study of high-molecular-weight proteins by solution NMR spectroscopy. Nat Protoc 1:749–754. doi:10.1038/nprot.2006.101

Tugarinov V, Kay LE. 2005. Methyl groups as probes of structure and dynamics in NMR studies of high-molecular-weight proteins. ChemBioChem 6:1567–1577. doi:10.1002/cbic.200500110

Vallat B, Madrid-Aliste C, Fiser A. 2015. Modularity of Protein Folds as a Tool for Template-Free Modeling of Structures. PLOS Comput Biol 11:e1004419. doi:10.1371/journal.pcbi.1004419

Van Der Spoel D, Lindahl E, Hess B, Groenhof G, Mark AE, Berendsen HJC. 2005. GROMACS: Fast, flexible, and free. J Comput Chem 26:1701–1718. doi:10.1002/jcc.20291

Webb B, Sali A. 2014. Comparative Protein Structure Modeling Using MODELLER. Curr Protoc Bioinforma 47:5.6.1-5.6.32. doi:10.1002/0471250953.bi0506s47

Wuthrich K. 1986. NMR of proteins and nucleic acids. Georg Fish Bak non-resident Lecturesh Chem Cornell Unversity.

