## Supplemental Images for "Protein Fold Determination by Assembling Extended Super-Secondary Structure Motifs Using Limited NMR Data"

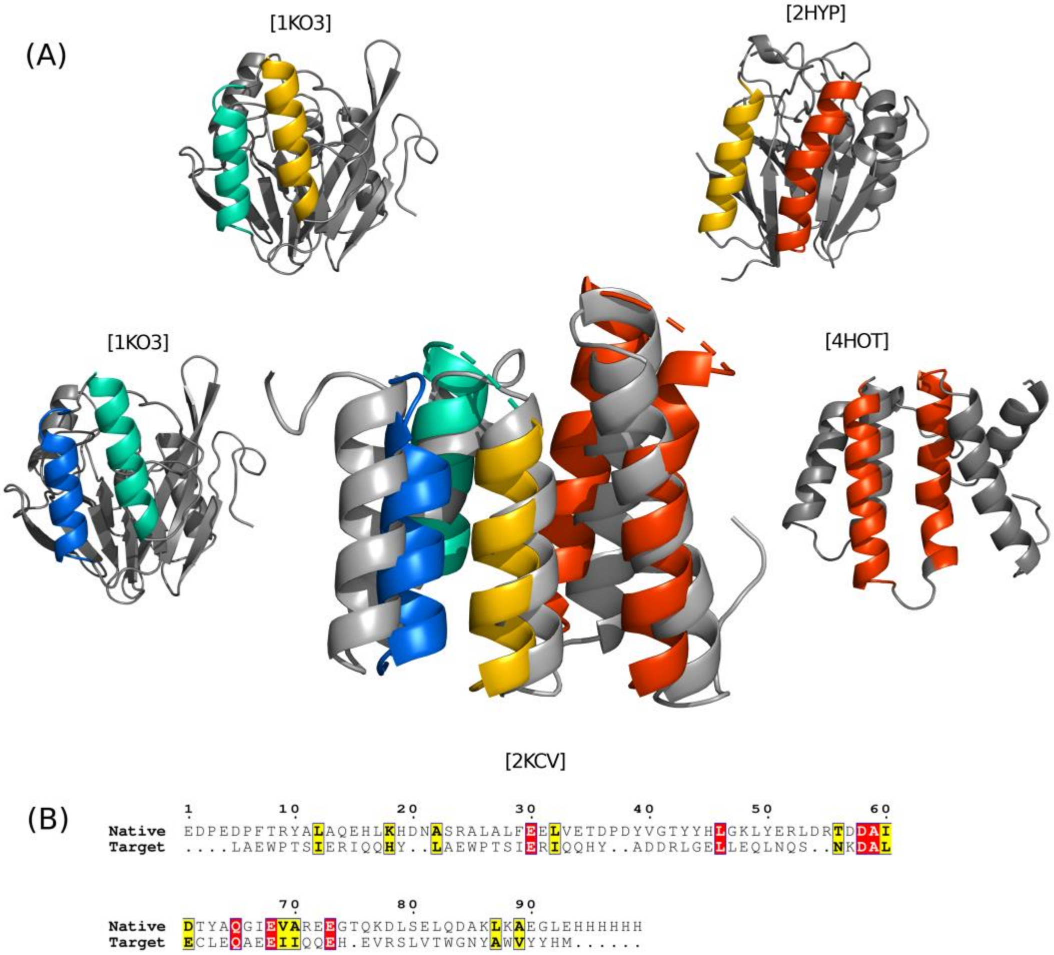


**Figure S1:** The Smotif assembly performed by DINGO-NOE-RDC algorithm on the protein Srr115c, Target-C [PDBID:2KCV]. (A) Cartoon representations of the final model superimposed onto the crystal structure (in gray) and of the parent proteins from which the Smotifs were derived. The respective parent proteins are labeled with their PDBID. The corresponding coloring highlights the Smotifs of the parent proteins and in the final structure. The target protein contains two tetratricopeptide repeat (TPR) domains spanning first four N-terminal helices (blue, green, yellow and red). The Smotif from 1KO3 were repeated twice to assemble the first three N-terminal helices of the target. The first and second helix of the 1KO3 (blue and green) are also equivalent and have a $C^{\alpha}$ RMSD of 0.1Å when the first helix is superimposed onto the second helix. The fourth helix is three residues longer than the first three helices and is not covered by the Smotif from 1KO3. (B) Sequence alignment between the target's native sequence and assembled Smotifs is shown. Target-C's sequence was generated by concatenating the individual sequences from the assembled Smotifs. Identical amino acid residues are highlighted in red and similar residues are highlighted in yellow. The sequence alignment is generated by placing equivalent residues between the target and the assembled Smotifs through secondary structure assignment.

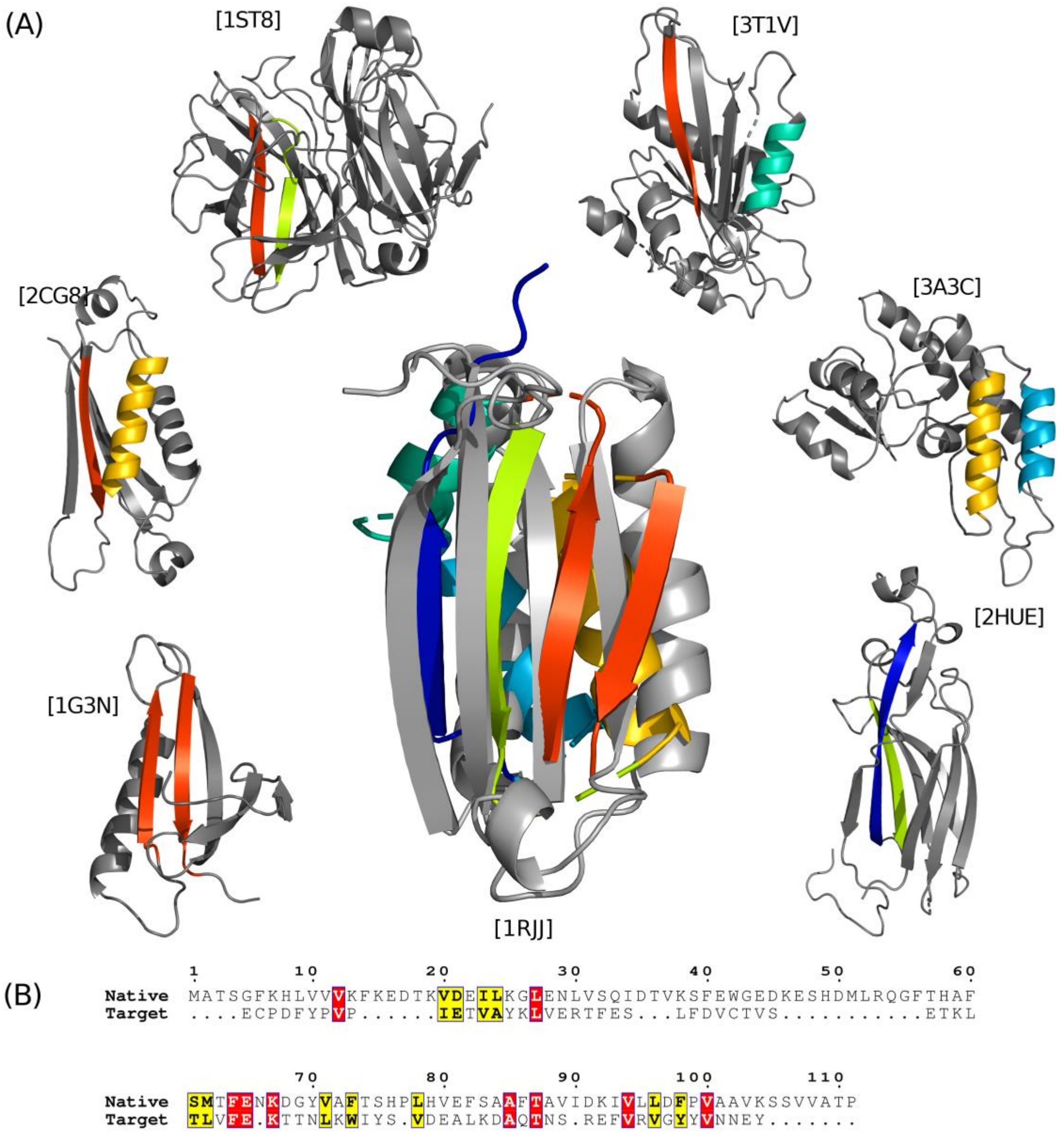


**Figure S2:** The Smotif assembly performed by DINGO-NOE-RDC algorithm on the target protein At5g22580 (Target-D) [PDBID:1RJJ]. (A) Cartoon representations of the final model superimposed onto the crystal structure (in gray) and of the parent proteins from which the Smotifs were derived. The respective parent proteins are labeled with their PDBID. The corresponding coloring highlights the Smotifs of the parent proteins and in the final structure. (B) All-atom model of the Smotif assembled structure from (A) generated using comparative modeling algorithm.

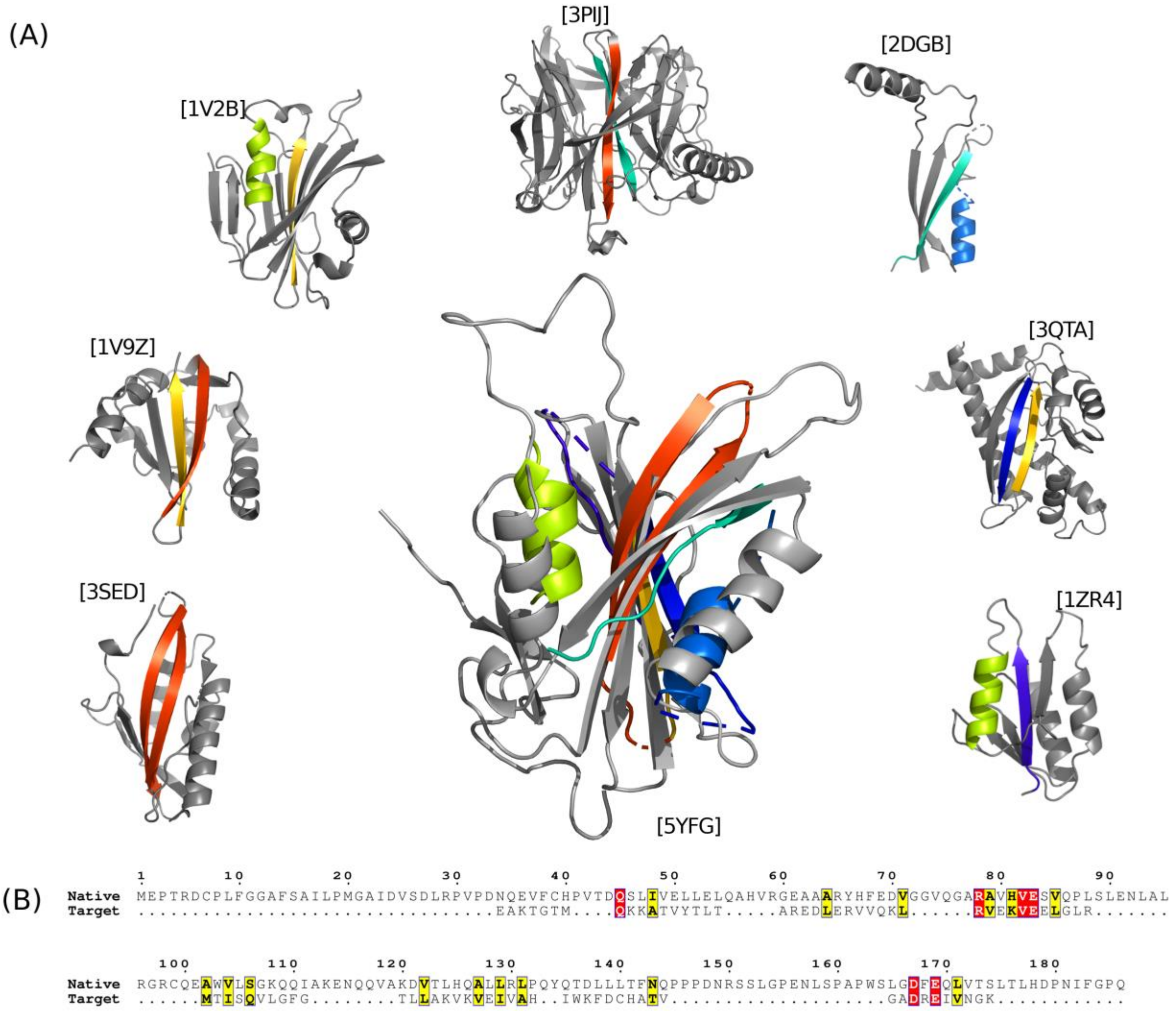


**Figure S3:** The Smotif assembly performed by DINGO-NOE-RDC algorithm for target protein Human MOG1 (Target-E). The panel descriptions are same as described in Figure S2 except the reference NMR structure used is PDBID:5YFG

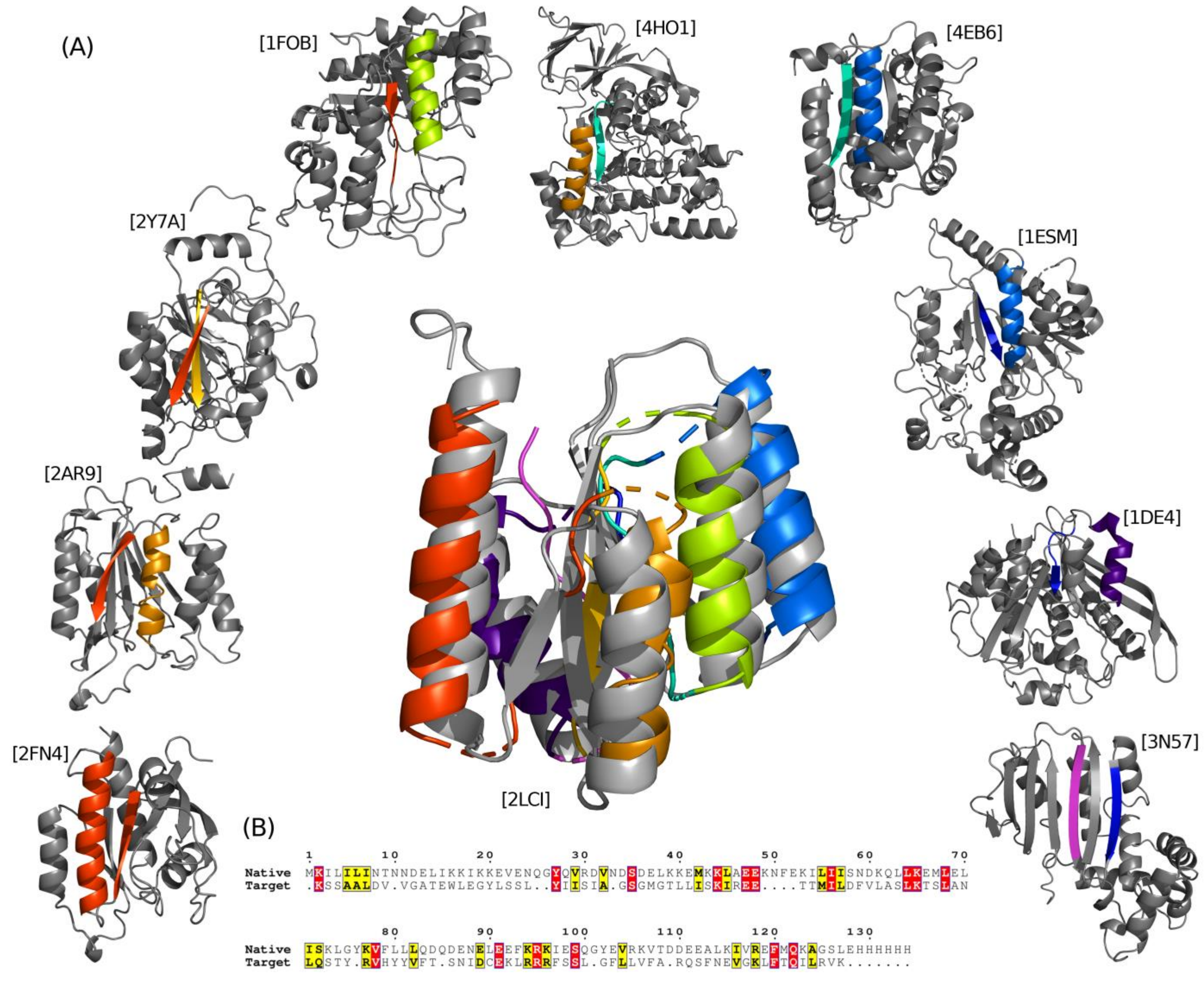


**Figure S4:** The Smotif assembly performed by DINGO-NOE-RDC algorithm for a *De novo* designed protein (Target-F). The panel descriptions are same as described in Figure S2 except the reference NMR structure used is PDBID:2LCI

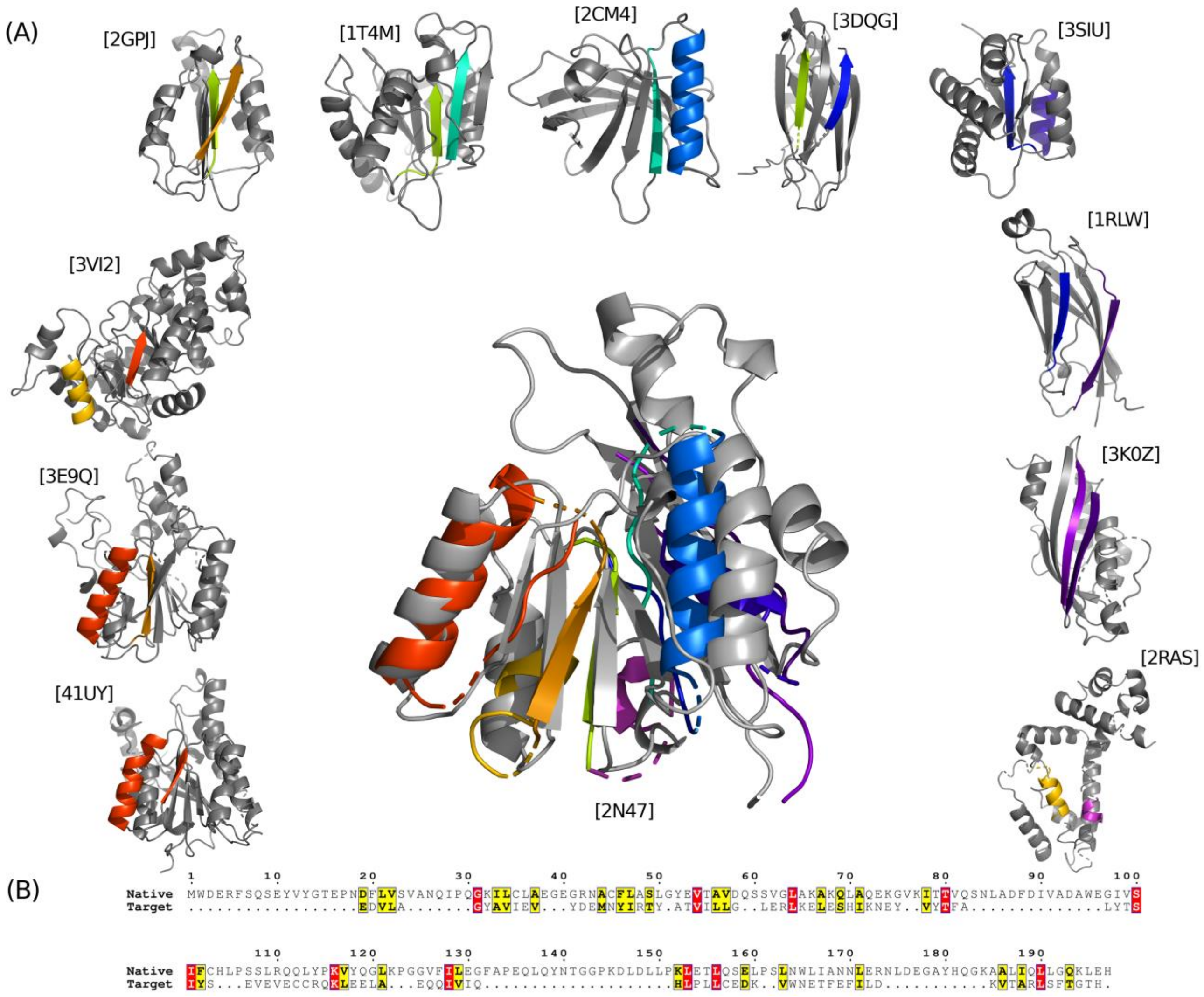


**Figure S5:** The Smotif assembly performed by DINGO-NOE-RDC algorithm for the target protein SgR145 (Target-G). The panel descriptions are same as described in Figure S2 except the reference NMR structure used is PDBID:2N47

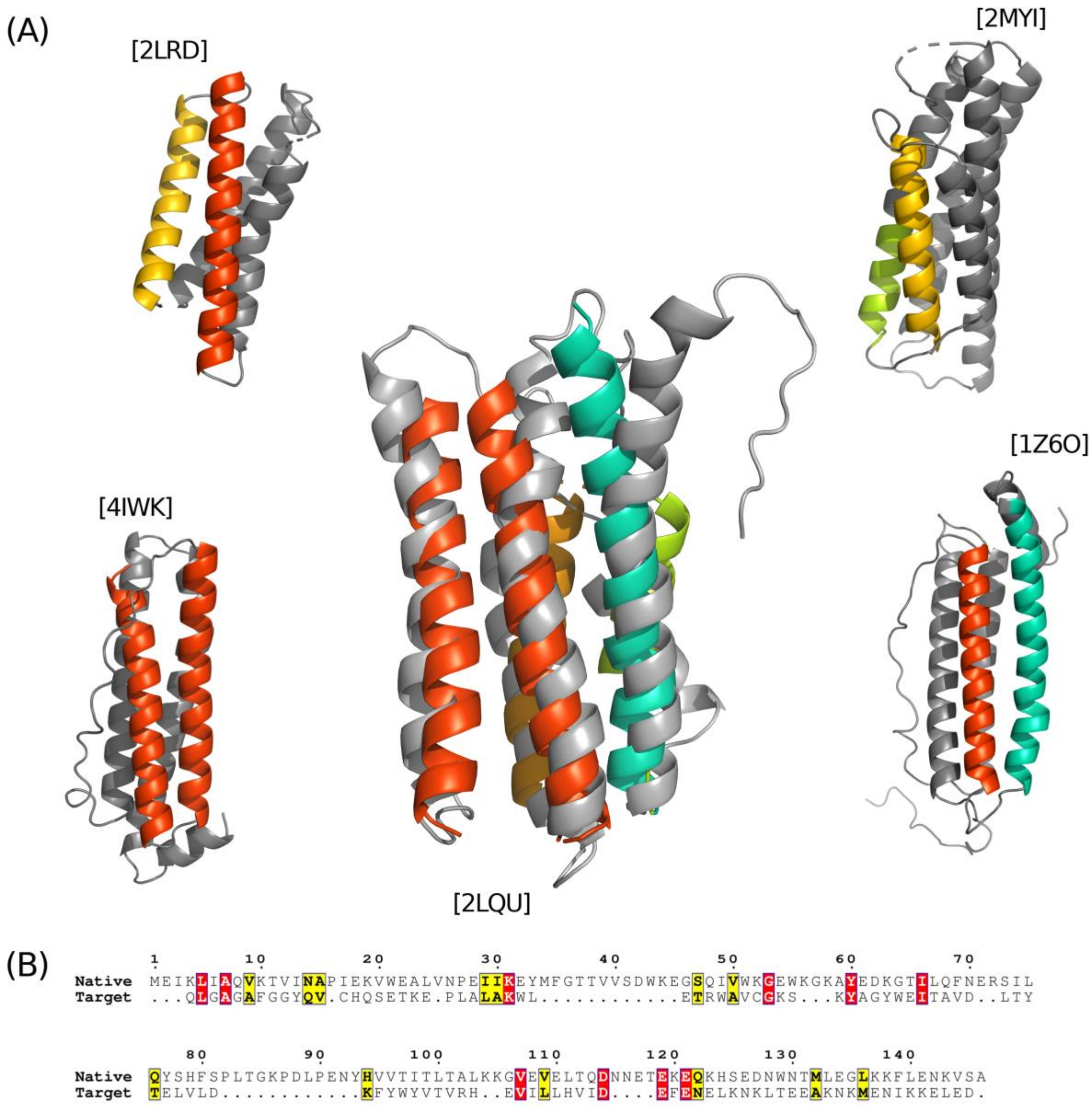


**Figure S6:** The Smotif assembly performed by DINGO-NOE-RDC algorithm for the target protein Decorbin binding protein (Target-H). The panel descriptions are same as described in Figure S2 except the reference NMR structure used is PDBID:2LQU

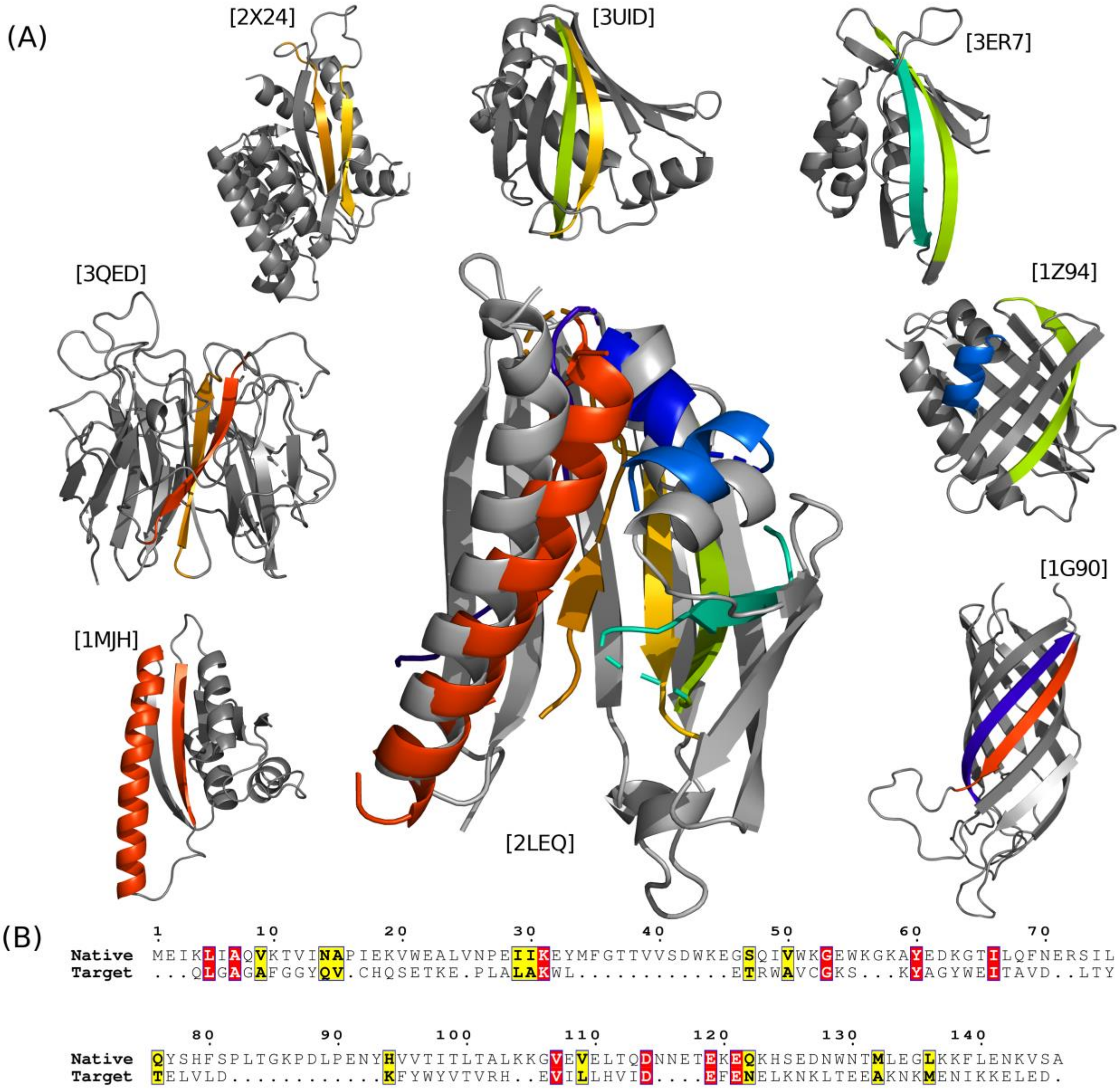


**Figure S7:** The Smotif assembly performed by DINGO-NOE-RDC algorithm for the target protein ChR145 (Target-I). The panel descriptions are same as described in Figure S2 except the reference NMR structure used is PDBID:2LEQ

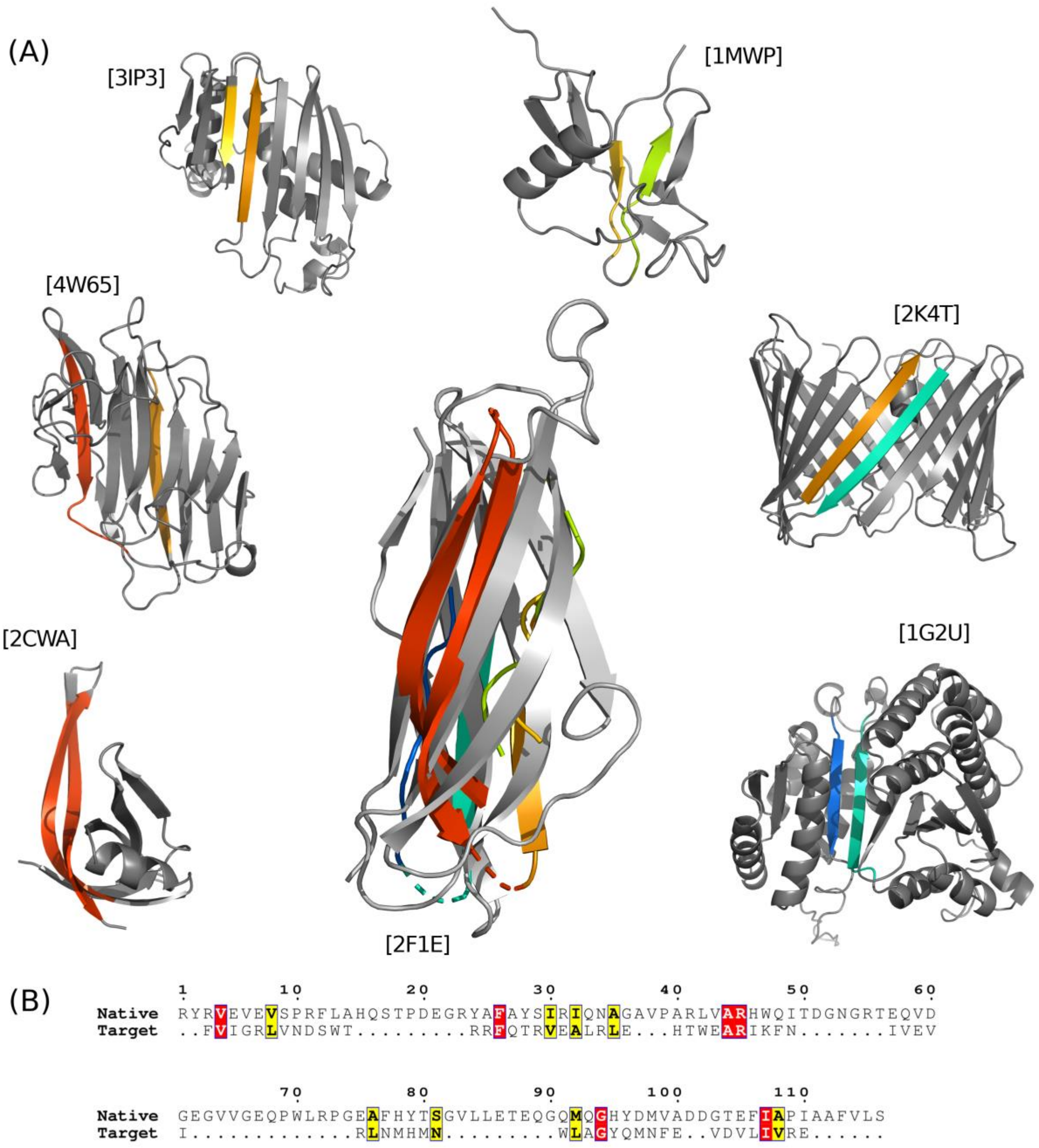


**Figure S8:** The Smotif assembly performed by DINGO-NOE-RDC algorithm for the target protein ApaG (Target-J). The panel descriptions are same as described in Figure S2 except the reference NMR structure used is PDBID:2F1E

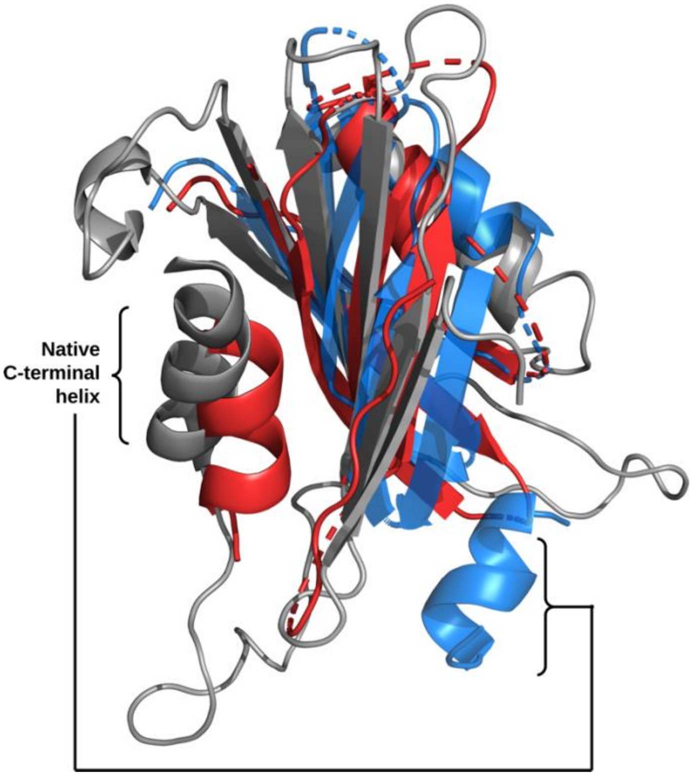


**Figure S9:** Degenerate Smotif assemblies for Target-E. Two top scoring Smotif assemblies are shown in red and blue. Both Smotif assemblies have similar NOE and RDC fit score. There were no NOEs between the C-terminal helix to the rest of the protein and only one RDC dataset which places the C-terminal helix in native (red) and non-native (blue) conformations.
